# Novel Partitivirus Infection of Bat White-nose Syndrome (WNS) Fungal Pathogen *Pseudogymnoascus destructans* Links Eurasian and North American Isolates

**DOI:** 10.1101/059709

**Authors:** Ping Ren, Sunanda S. Rajkumar, Haixin Sui, Paul S. Masters, Natalia Martinkova, Alena Kubátová, Jiri Pikula, Sudha Chaturvedi, Vishnu Chaturvedi

**Author notes:** Current address: University of Texas Medical Branch, Galveston, TX, USA. Current address: ICMR Medical Research Institute, Pudicherry, India.

## Abstract

Bat White-nose Syndrome (WNS) fungus *Pseudogymnoascus destructans* had caused mass mortality in the North American bats. A single clone of the pathogen (Hap_1) was likely introduced in the United States while Eurasian population comprised of several haplotypes. The origin and spread of *P. destructans* remain enigmatic due in part to a lack of precise population markers. We searched for *P. destructans* mycoviruses as they are highly host-specific, and their spread could provide a window on the origin of the host fungus. We discovered a *P. destructans* bipartite virus PdPV-1 with two double-stranded RNA (dsRNA) segments - LS (1,683 bp) and SS (1,524 bp) with motifs similar to viral RNA-dependent RNA polymerase (RdRp) and putative capsid proteins (CPs), respectively. Both LS and SS ORFs were embedded only in the positive strand of each dsRNA segment. Sequence alignments and phylogenetic analysis suggested that both segments constitute the genome of a new virus similar to the mycoviruses in the family *Partitiviridae* genus *Gammapartitivirus.* Purified viral particles appeared as isometric virions with approximately 33 nm diameters typical of partitiviruses. A newly developed RT-PCR assay revealed that all US isolates and only a few Eurasian isolates were infected with PdPV-1. PdPV-1 was *P. destructans* - specific as closely related non-pathogenic fungi *P. appendiculatus* and *P. roseus* tested negative. Thus, PdPV-1 establishes a link between the Eurasian and North American *P. destructans.* PdPV-1 could be used as an experimental tool to further investigate fungal biogeography, and the host - pathogen interactions.

## IMPORTANCE

The fungal disease White-nose Syndrome threatens many bat species in North America. The pathogen *Pseudogymnoascus destructans* appeared first in Upstate New York nearly a decade ago. The population markers used to date revealed that US population of *P. destructans* comprised of a single clone (Hap_1) while the Eurasian population was diverse. More precise tools are still needed to pinpoint the exact source of this devastating disease. We studied *P. destructans* for viral infections (mycoviruses). Fungal viruses are highly host specific, and their spread mirrors the distribution of the host fungus. We found a double-stranded RNA virus (PdPV-1) with size, shape and sequences similar to fungal partitiviruses. All the US *P. destructans* and a few Eurasian isolates were positive for PdPV-1. Other closely related *Pseudogymnoascus* species also found in the bat hibernacula, were free of PdPV-1 infection. These findings provide a new avenue to study *P. destructans* origin and pathogenic attributes.

*Pseudogymnoascus destructans* is a psychrophilic (cold-loving) fungus responsible for bat White-nose Syndrome (WNS) (1– 3). *Pseudogymnoascus destructans* infects at least eleven species of bats in North America, of which seven species exhibited disease symptoms. The infected bat species include the endangered Indiana bat (*Myotis sodalist*) and gray bat (*M. grisescens*) (https://www.whitenosesyndrome.org/about/bats-affected-wns). Several species including widely distributed little brown bats (*M. lucifugus*) face local extirpations (4). *Pseudogymnoascus destructans* infects bats during hibernation in the caves and abandoned mines. The fungus also persists in the environment in the absence of bats (5). *Pseudogymnoascus destructans* isolates analyzed to date from the US, and Canada represented a clonal population (6, 7). Although European bats also suffer from *P. destructans* infections, the pathogen population comprise of many haplotypes including Hap_1, which is the only haplotype found in fungal isolates from North America (8, 9). Despite apparent genetic homogeneity, phenotypic variations were reported among North American *P. destructans* isolates specifically in the mycelial growth rate, exudate production, and pigment formation and diffusion into agar media (Figure 1A) (7). The underlying cause (s) for the phenotypic variations remain to be discovered.

**Fig. 1A.**
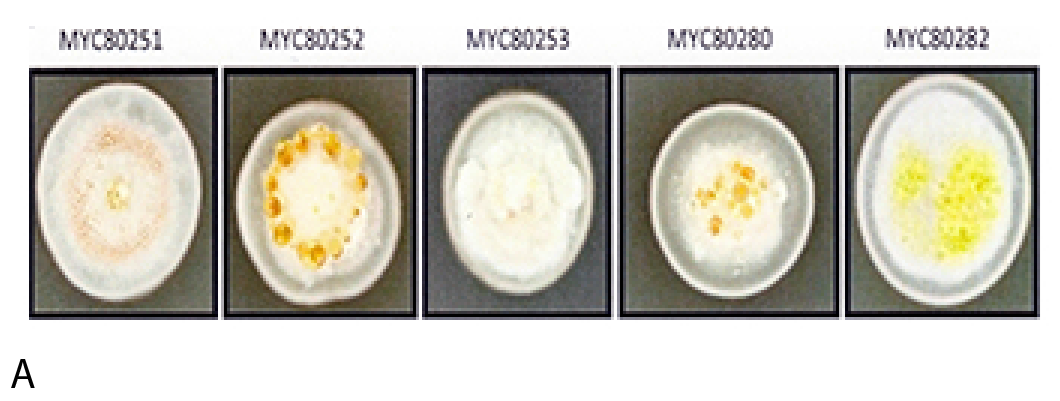
Phenotypic variation in clonal population of *P. destructans*. There is notable variation of colony color, texture, exudates and growth rates among the US, and Canadian isolates.

Virus infections in fungi (‵mycovirus′) are rather common and latent mostly (10, 11). The majority of mycoviruses are dsRNA isometric particles although ssRNA, and ssDNA mycoviruses have been recognized. Three well-studied mycovirus systems include: i) yeast killer toxins in *Saccharomyces cerevisiae* and non-conventional yeasts (12), ii) hypovirulence in *Cryphonectria parasitica* and *Ophiostoma ulmi*, the etiologic agents of Chestnut blight and Dutch Elm disease, respectively (13, 14), and iii), symbiotic role in fungal endophytes of grasses (15). There are reports linking mycovirus infection to the phenotypic changes in the fungal host (16).

Recently dsRNA mycoviruses in human pathogen *Aspergillus fumigatus* were found to cause aberrant phenotypes and hypervirulence(17). Mycoviruses could also provide a ‵phylogenomics window′ as their evolution showed strong a co-divergence with their fungal hosts (18). We hypothesized that origin, evolution, and virulence of *P. destructans* could be investigated by focusing on the mycoviruses. The current study summarizes the molecular characterization of novel virus (named as PdPV-1) infection in *P. destructans.* All US and some Eurasian isolates tested positive for PdPV-1. PdPV-1 was host-specific to *P. destructans* as closely related non-pathogenic fungi *P. appendiculatus* and *P. roseus* tested negative for the mycovirus (3).
The results suggest that PdPV-1 links the Eurasian and North American *P. destructans,* and provides an experimental tool for the future investigations of host - pathogen interactions (Figure 1).

Multiple bands were observed on the agarose gel in total nucleic acids extracted from *P. destructans* (MYC80251) (Figure 1B, lane 1). Treatment of nucleic acids extracts with various nucleases and DNase I yielded double bands thereby, suggesting the presence of dsRNA (Figure 1B, lane 4). dsRNA bands were estimated to be 1.5 – 2.0 kb (Figure 1B, bottom panel). Two dsRNA segments- LS and SS were identified from sequence analysis. The LS segment contained an open reading frame (ORF) that encodes 539 amino acids with a molecular mass of approximately 62.7 kDa. The small segment SS comprised 1,524 bp nucleotides with 49% GC content. The SS segment contains an ORF that encodes 434 amino acids with a molecular mass of approximately 46.9 kDa. Both ORFs were identified only on the positive strand of each dsRNA segment. The negative strands did not contain any significant ORFs that are longer than 82 amino acids. The 5' untranslated regions of the plus strands of segments LS and SS were 8 and 60 nucleotides, respectively; whereas the 3' untranslated regions of the plus strands of segments LS and SS were 54 and 159 nucleotides, respectively. GenBank search revealed that the ORFs of dsRNA LS and SS have significant similarities to the putative RNA-dependent RNA polymerase (RdRp) and the capsid protein (CP), respectively, of viruses from the family *Partitiviridae* genus *Gammapartitivirus.* The partitivirus was named *Pseudogymnoascus destructans* partitivirus (PdPV-1). Phylogenetic trees derived from both RdRp and CP sequences exhibited three major branches and supported earlier conclusion that PdPV-1 is a member of the genus Gammaartitivirus, the family Partitiviridae (Figures 1 and supplement). Also, phylogenetic analyses of the putative RdRp and CP of PdPV-1 showed that PdPV-1 was much closely related to the gammapartitiviruses of PsV-S. Transmission electron microscopy (TEM) confirmed the existence of viral particles in *P. destructans*. Several isometric PdPV-1 virions with estimated diameter of about 33 nm, characteristics of partitiviruses, were observed (Figure 1C). A specific RT-PCR revealed that all *P. destructans* isolates from the US were positive for PdPV-1. A few Eurasian strains also tested positive for PdPV-1. The closely related fungi *P. roseus* and *P. appendiculatus* did not yield any detectable LS or SS segment of PdPV-1 dsRNA (Table S1).

**Fig. 1B.**
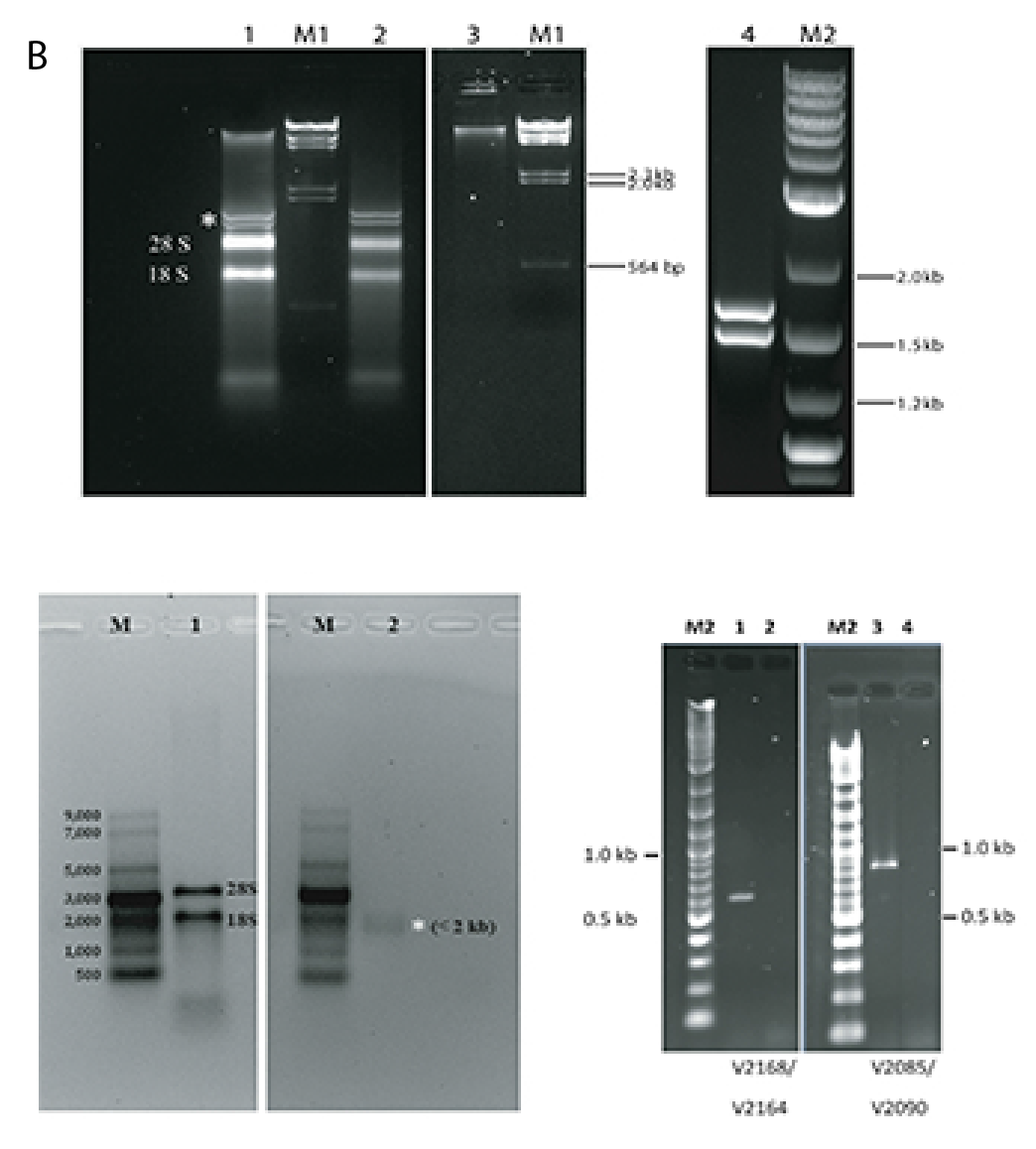
*P. destructans* nucleic acid extract analysis. An analytical run on 1% native agarose gel showed bands suggestive of genomic DNA, ribosomal RNA and two additional discrete nucleic acid bands (asterisk).These bands were determined to be RNA species since RNase, but not DNaseI digestion, led to their disapperance. Lane M1: λ-Hind III marker. Lane M2: 2-LOG DNA marker. Lane1: Total nucleic acid. Lane 2: Total nucleic acid + DNase I. Lane 3: Total nucleic acid + RNase III + RNase A. Lane 4: Total nucleic acid + S1 Nuclease + DNase I. Size of the RNA bands. RNA were sized by an analytical run of the purified samples on denaturing polyacrylamide gel The size of the two RNA bands is less than 2 kb. These RNA bands were maked by the 18S ribosomal RNA band in total RNA samples. Lane M: ssRNA marker (in bp). Lane 1: Total RNA. Lane 2: Gel purified particle.

**Fig. 1C.**
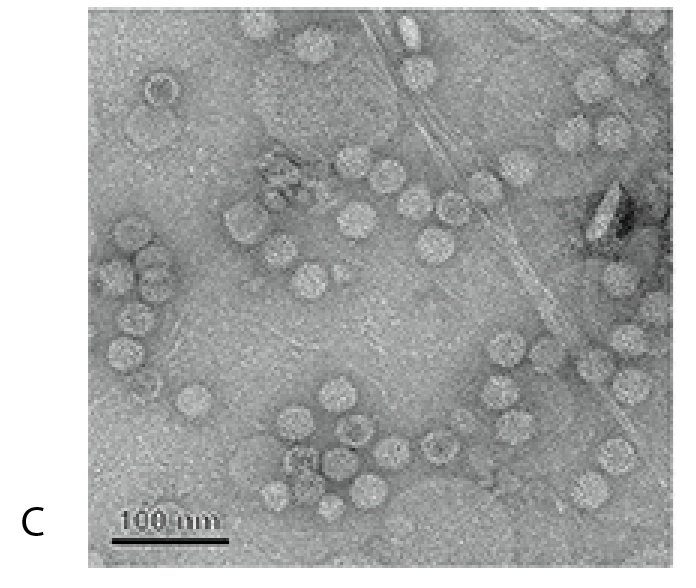
Transmission electron microscopy of purified PdPV-1. Several isometric viral particles are visible approximately 33 nm in diameter. The bar represents 100 nm.

**Fig. 1D.**
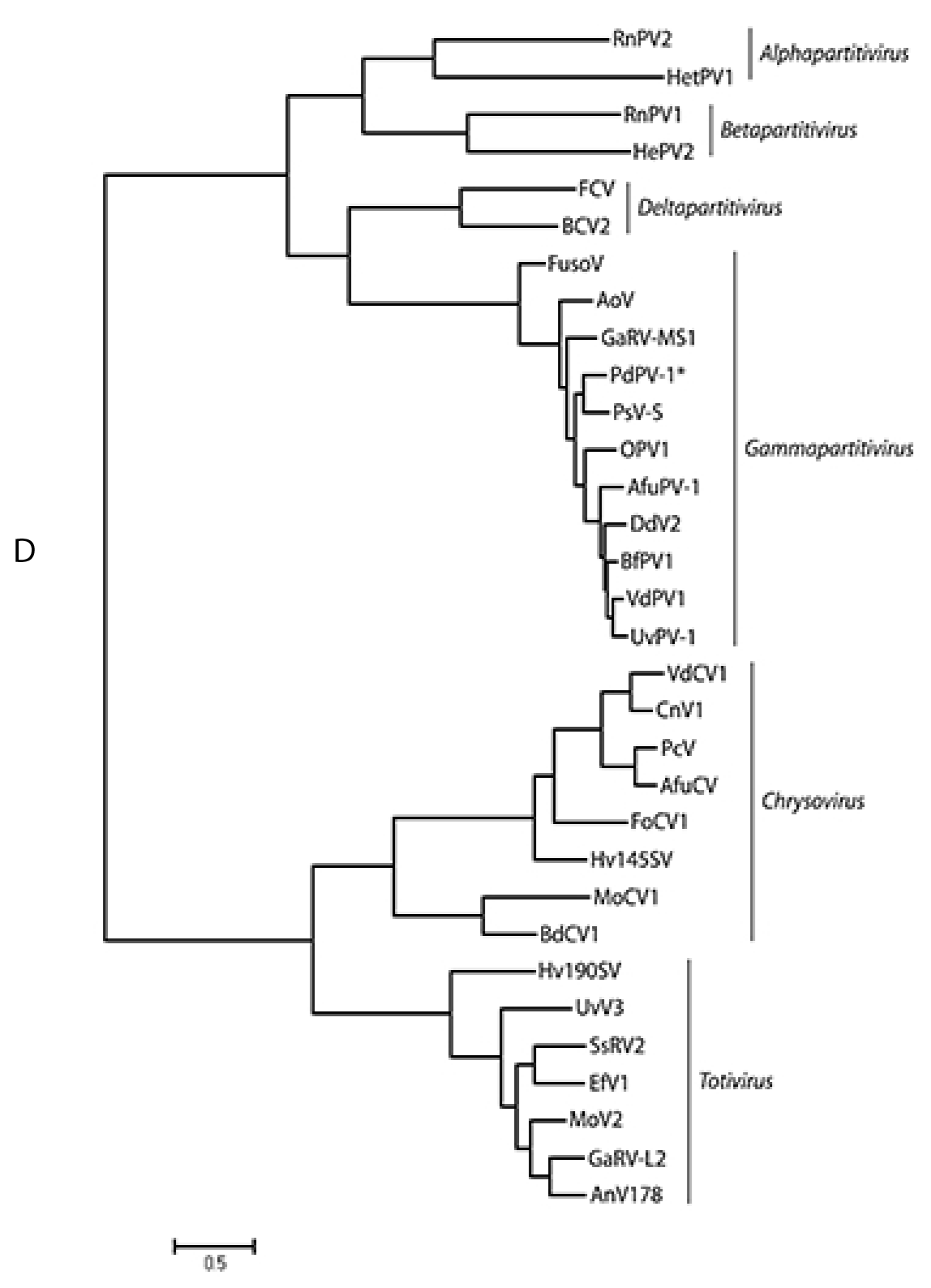
Phylogenetic analyses of PdPV-1. RdRp amino acid sequences of the representative members of the family *Partitiviridae, Totiviridae* and *Chrysoviridae* were used to construct Maximum likelihood phylogenetic tree with MEGA 6 (GenBank accession numbers are provided in Table S3).

We confirmed the discovery of *P. destrucatns* mycovirus PdPV-1 by:(i) demonstration of dsRNA by DNA and RNA endonucleases, (ii) nucleotide similarities of the cloned dsRNA viral genome, and their phylogenetic alignments with mycoviruses from the family *Partitiviridae* genus *Gammapartitivirus*, (iii) TEM confirmation of isometric viral particles, (iv) high degree of host specificity for *P. destructans* and absence from the closely related *Pseudogymnoascus* species, and (v) exclusive association of PdPV-1 with *P. destructans* Haplotype 1 (Hpa_1), which is the clonal genotype of all North American isolates and only found in a select number of Eurasian isolates (6, 8, 9). Overall, PdPV-1 showed size, shape and nucleotide sequences typical of the fungal viruses in the family *Partitiviridae* genus *Gammmapartitivirus* (19)

The discovery of dsRNA mycovirus in *P. destructans* is consistent with the wide occurrence of mycoviruses in fungi (10, 20). Earlier observations suggested that partitiviruses caused cryptic infections in their hosts without any discernible phenotypic changes (10, 16). A more evolved pattern was discerned in recent studies with viral induced hypovirulence in plant pathogenic fungi without visible phenotypic effects; virus infections of animal pathogenic fungi and protozoan pathogens showed hypervirulence (11, 21, 22). Although our limited testing was inconclusive, in-depth experiments are needed to discern the outcome of PdPV-1 infection on *P. destructans* phenotype and virulence. The absence of PdPV-1 infection in other *Pseudogymnoascus* species, collected from the same bat hibernacula in the US, suggested a close viral association with *P. destructans*. The observed host-specificity of PdPV-1 was consistent with the narrow host ranges of mycoviruses due to severe bottlenecks in their horizontal transmission (11).

The PdPV-1 discovery provided an independent and corroborative evidence for the emergence of *P. destructans* as a novel pathogen (8, 23). The local origin of PdPV-1 in the US was unlikely due to uniform infection of *P. destructans* isolates tested so far as well as the failure of viral-curing experiments (supplementary information). A recent Eurasian origin was the most likely explanation based upon the findings of both viral-free and viral-infected *P. destructans* isolates in the Eurasian populations (Figure 1E). In such a scenario, the likely introduction of PdPV-1 to the US took place by the arrival of a mycovirus infected isolate of *P. destructans* (Hap_1 haplotype). Presently, the influence of different haplotypes of *P. destructans* on fungal virulence and other attributes remains unknown (9). The association between PdPV-1 and Hap_1 could become crucial if Hap_1 represented a more virulent pathotype of *P. destructans* (24)

PdPV-1 LS and SS nucleotide sequences are available from NCBI GenBank. (Accession number: KP128044 and KP128045, respectively)

**Fig. 1E.**
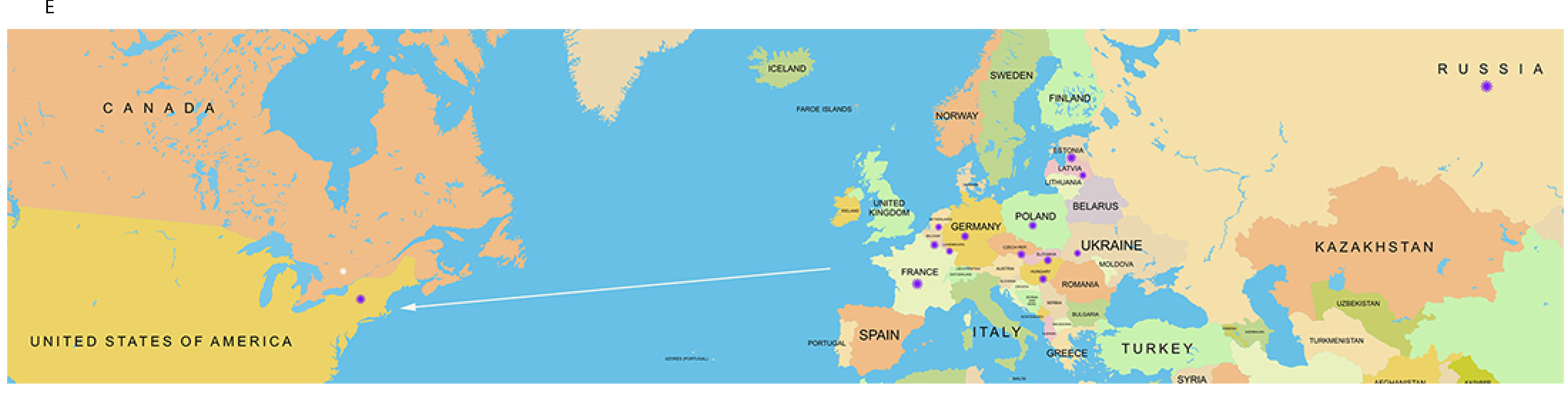
Projected origin and transmission of *P. destructans*. The country map highlights areas where PdPV-1 infected fungal isolates were found. Eurasian samples also contained viral negative samples.

## ACKNOWLEDGEMENTS

This study was funded in parts with the US Fish & Wildlife Service (F15AP00948) and the National Science Foundation (1203528) research awards to VC and SC. DNA sequencing was performed at the Wadsworth Center Molecular Genetics Core.

## Supplementary Information

<u>Fungal isolates</u>. *P. destructans* isolate (MYC80251) and other US and Eurasian isolates have been described previously (1– 3). The isolates from other *Pseudogymnoascus* species were also tested (4). The supplementary table 1 provides additional details of test strain. All fungal isolates were routinely maintained on Sabouaraud Dextroxe Agar or Potato Dextrose Agar at 15°C. The cultures were stored in 15% glycerol at -80°C. The methods for the observations of colony morphology and extra-cellular enzymes was described previously (1, 5).

<u>dsRNA extraction</u>. *P. destructans* was grown in a stationary culture in potato dextrose agar (PDA) at 15°C for 2-3 months. About 1 g wet weight mycelium was harvested and grounded to powder in the presence of liquid nitrogen with mortar and pestle. The powder was collected and suspended in 1 ml extraction buffer (150 mM sodium acetate, pH 5.0, 100 mM LiCl, 4% sodium dodecyl sulfate, 10 mM EDTA, pH 8.0, and 20 mM β-mercaptoethanol) and incubated on ice for 10 min. Total nucleic acids were obtained by traditional phenol-chloroform extraction and LiCl, 2-propanol precipitation method. ssRNA and DNA were removed by treatment with S1 nuclease and DNase I (Life Technologies, Carlsbad, CA). The extractions were run on 1% agarose gel, stained with ethidium bromide and visualized with UV transillumination. To confirm the dsRNA nature of the mycovirus, the total nucleic acid solutions were treated with RNase III and RNase A (Life Technologies) to see the disappearance of dsRNA. Denaturing polyacrylamide gel was used to run the purified dsRNA extractions to see the size of two RNA bands (6, 7).

<u>cDNA synthesis and sequence analysis</u>. Purified dsRNA fractions containing 2 segments were denatured in 90% dimethyl sulfoxide (DMSO) at 65°C for 20 min in the presence of random hexadeixinucleotide and quickly chilled on ice. Then first-strand cDNA was synthesized using M-MLV reverse transcriptase (Life Technologies). The resulting cDNA were amplified with random primers and the variously sized PCR products were cloned in TOPO TA cloning vector. A series of overlapping cDNA clones were obtained from the sequencing of the positive clones. Determination of the ends of each dsRNA was done using FirstChoice^®^ RLM-RACE Kit (Life Technologies) (8).

<u>Nucleotide sequence analysis</u>. The nucleotide sequence contigs were assembled by Sequencher 4.8 (Gene Codes Co., Ann Arbor, MI). The conserved sequences were BLAST searched in GenBank by tblastx program. Multiple sequence alignments for two dsRNA segments were carried out using ClustalW analysis by MacVector 7.2 (Accelrys Inc., Cary, NC) respectively. The phylogenetic analyses were conducted using maximum likelihood method by MEGA 6.0 (9).

<u>Virus purification</u>. About 60 g wet weight fungal mycelium were collected and grounded to powder as described in dsRNA extraction session. The homogenates were mixed with extraction buffer (0.1 M sodium phosphate, pH 7.4, containing 0.1% β-mercaptoethanol) at a volume of 5 ml/g of wet mycelium. The suspensions were vortex hard after adding equal volume of chloroform and broke the emulsion by centrifuging in a Sorvall GSA rotor at 8,000 rpm for 20 min. The upper aqueous layer was then mixed thoroughly with 0.5 volumes of 30% polyethylene glycol 8000 (PEG) in 0.85% NaCl and hold on ice for 1 hour. PEG precipitates were pelleted in a Sorvall GSA rotor with 16,000× g at 4°C for 30 min and resuspended in 0.1 M sodium phosphate, pH 7.4. After centrifuge with 23,000× g at 4°C for 20 min to remove unsuspended debris, the supernatant was ultra-centrifuged in a Beckman SW42 rotor at 76,000× g, 4°C for 2 hrs to pellet the virus. The pelleted virus was re-suspended in a total of 4 ml 0.1 M sodium phosphate buffer and purified by loading the viral suspension onto pre-formed gradients of 10% - 50% (w/v) sucrose in 0.1 M sodium phosphate buffer and centrifuging at 76,000× g, 4°C overnight. The collected virus fractions were diluted in 0.1 M sodium phosphate buffer and preserved at 4°C for immediate electron microscopy (10– 12).

<u>Transmission electron microscopy</u>. Two μl of the virus solution was placed on a glow-discharged copper grid covered with a continuous carbon film. After 1 minute of adsorption, the grid was washed with pure water for several seconds and stained with 3μl of 2% (w/v) uranyl acetate solution for 1 minute. The staining solution was blotted away by Whatman No. 1 filter paper. The grid was air dried completely before it was examined in a JEOL JEM-1400 electron microscope operating at 120 keV (13). The micrographs were recorded at various magnifications using a 4K × 4K CMOS camera (TVIPS F-416).

<u>Reverse transcript PCR (RT-PCR) assay</u>. First-strand cDNA synthesis was performed as described earlier and followed by PCR using primer pairs V2085/V2090 and V2168/V2164 to amplify SS and LS segments respectively. Then the PCR products were sequenced using the same primers and aligned to confirm the sequences.

<u>Viral curing</u>. To cure the viruses from *P. destructans* isolates, tiny hyphal tips of isolate MYC80251 were inoculated on PDA with 5 μg/ml cycloheximide (14) and limited nutrient cAMP-rifamycin agar plate (15). After 7 days incubation at 15°C, when there was a discernible increase in the colony growth, hyphal tips were cut with sterile blade and transferred on the fresh PDA - cycloheximide plates and cAMP - rifamycin agar plates. This passage was repeated for five times. For the last passage, the fungal colonies were allowed to grow for about 10 days to obtain sufficient hyphal mass for total RNA extraction and the presence of mycoviruses examined by RT-PCR.

<u>Detection of PdPV-1 dsRNAs or virus particles in broth culture supernatant</u>. *P. destructans* isolates MY80251, PSEU14, and LBB17 were cultured in a 125 ml flask containing 25 ml Potato dextrose broth medium at 15°C without shaking for 4 weeks. Five ml of culture supernatants were centrifuged at 17,000 g for 5 min to remove conidia and free cells. Then 250 μl resultant supernatants were used for nucleic acids extraction with SDS/phenol and subjected to agarose gel electrophoresis. To detect virus particles, 4 ml resultant supernatants were ultracentrifuged at 148,400 g for 1 hour and the precipitate was suspended in 0.05 M Tris/HCl (pH 7.8) containing 0.15 M NaCl at 4°C. The suspension was stained and visualized by transmission electron microscope as described earlier.

**Supplementary Table 1.** Fungal isolates used in this study

| *Pseudogymnoascus destructans* | Location | Sources | Viral RNA |
| --- | --- | --- | --- |
| MYC80251 | Albany, New York | *Myotis lucifugus* | + |
| PESU14 | Avery, North Carolina | *Perimyotis subflavus* | + |
| LBB17 | Lawrence, Ohio | *Myotis lucifugus* | + |
| PSEU8 | Greenbrier, West Virginia | *Perimyotis subflavus* | + |
| LBB11 | Woodward, Pennsylvania | *Myotis lucifugus* | + |
| M2335 | Tompkins, New York |  | + |
| M2337 | Erie, New York |  | + |
| M2339 | Ulster, New York |  | + |
| HA-8-2 | Hailes Cave, New York | Cave wall swab | + |
| VTG1-5-1 | Greely Mine, Vermont | Soil | + |
| AC-3-3 | Aelous Cave, Vermont | Soil | + |
| *Pseudogymnoascus roseus* |  |  |  |
| 3-VT-5 | Vermont | Hibernacular soil | - |
| 5-NY-6 | New York | Hibernacular soil | - |
| 5-NY-8 | New York | Hibernacular soil | - |
| 5-NY-9 | New York | Hibernacular soil | - |
| WSF-3629 | Wisconsin | Amorphus peat | - |
| UAMH1658 | Canada | Sphagnum bog | - |
| *Pseudogymnoascus appendiculatus* |  |  |  |
| UAMH10510 | Canada | Wood bait block, Sphagnum bog | - |

**Supplementary Table 2.**
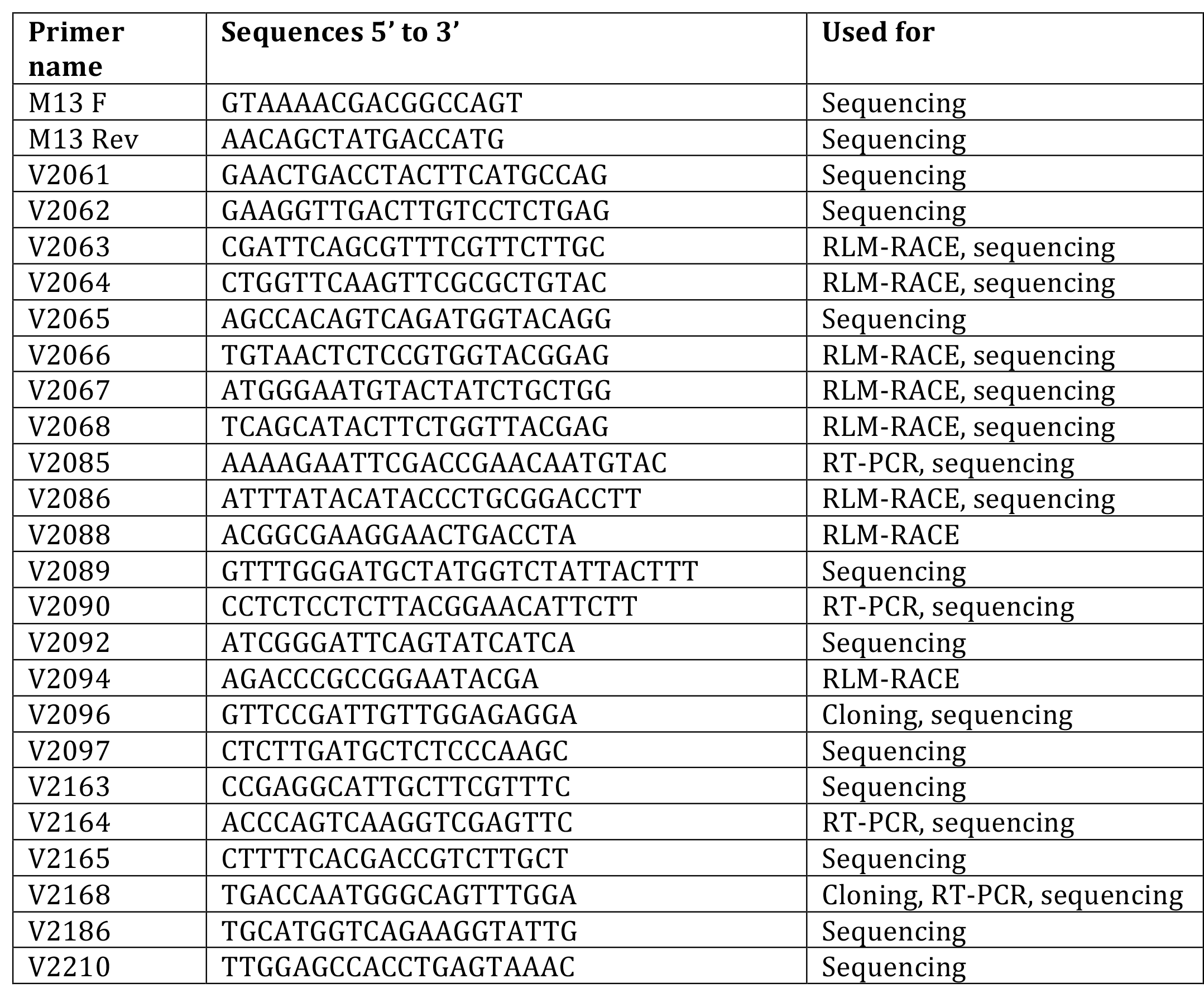
Details of primers used in this study.

**Supplementary Table 3.** Accession number and other details of amino acid sequences used for phylogenetic analyses.

| Mycovirus | Abbreviation | Amino acid sequences GenBank accession number |  |
| --- | --- | --- | --- |
|  |  | for RdRp | for Capsid |
| *Aspergillus fumigatus* chrysovirus | AfuCV | CAX48749 | CAX48751 |
| *Aspergillus fumigatus* partitivirus-1 | AfuPV-1 | CAY25801 | CAZ61323 |
| *Aspergillus mycovirus* 178 | AnV178 | ABX79995 | ABX79994 |
| *Aspergillus ochraceus* virus | AoV | ABV30675 | ABV30676 |
| *Beet cryptic* virus 2 | BCV2 | ADP24757 |  |
| *Botryosphaeria dothidea* chrysovirus 1 | BdCV1 | AGZ84312 | AGZ84313 |
| *Botryotinia fuckeliana* partitivirus 1 | BfPV1 | CAM33266 | CAM33267 |
| *Cryphonectria nitschkei* chrysovirus 1 | CnV1 | ACT79256 | ACT79252 |
| *Discula destructive* virus 2 | DdV2 | AAK59379 | AAK59380 |
| *Epichloe festucae* virus 1 | EfV1 | CAK02788 | CAK02787 |
| *Fig cryptic* virus | FCV | CBW77436 | CBW77437 |
| *Fusarium oxysporum* chrysovirus | FoCV1 | ABQ53134 | ABQ58816 |
| *Fusarium solani* mycovirus | FusoV | BAA09520 | BAA09521 |
| *Gremmeniella abietina* RNA virus L2 | GaRV-L2 | YP_044807 | YP_044806 |
| *Gremmeniella abietina* RNA virus MS1 | GaRV-MS1 | AII16004 | AII16002 |
| *Helminthosporium victoriae* 145S virus | Hv145SV | YP_052858 | YP_052859 |
| *Helminthosporium victoriae* virus 190S | Hv190SV | NP_619670 | NP_619669 |
| *Heterobasidion* partitivirus 1 | HetPV1 | ADV15441 | ADV15442 |
| *Heterobasidion* partitivirus 2 | HetPV2 | ADL66906 | ADL66905 |
| *Magnaporthe oryzae* chrysovirus 1 | MoCV1 | YP_003858286 | Not available |
| *Magnaporthe oryzae* virus 2 | MoV2 | YP_001649206 | YP_001649205 |
| *Ophiostoma* partitivirus 1 | OPV1 | CAJ31886 | CAJ31887 |
| *Penicillium chrysogenum* virus | PcV | YP_392482 | YP_392481 |
| *Penicillium stoloniferum* virus S | PsV-S | AAN86834 | AAN86835 |
| *Pseudogymnoascus destructans* virus 1 | PdV-1 | KP128044 | KP128045 |
| *Rosellinia necatrix* partitivirus 2 | RnPV1 | BAD98237 | BAD98238 |
| *Rosellinia necatrix* partitivirus 2 | RnPV2 | BAM78602 | BAK53192 |
| *Sphaeropsis sapinea* RNA virus 2 | SsRV2 | NP_047560 | NP_047559 |
| *Ustilaginoidea virens* partitivirus | UvPV-1 | AGO04402 | AGO04403 |
| *Ustilaginoidea virens* RNA virus 3 | UvV3 | YP_009004156 | YP_009004155 |
| *Verticillium dahliae* chrysovirus 1 | VdCV1 | ADG21213 | ADG21214 |
| *Verticillium dahliae* partitivirus 1 | VdPV1 | AGI52210 | AGI52209 |

## Supplementary figures

**Fig. S1.**
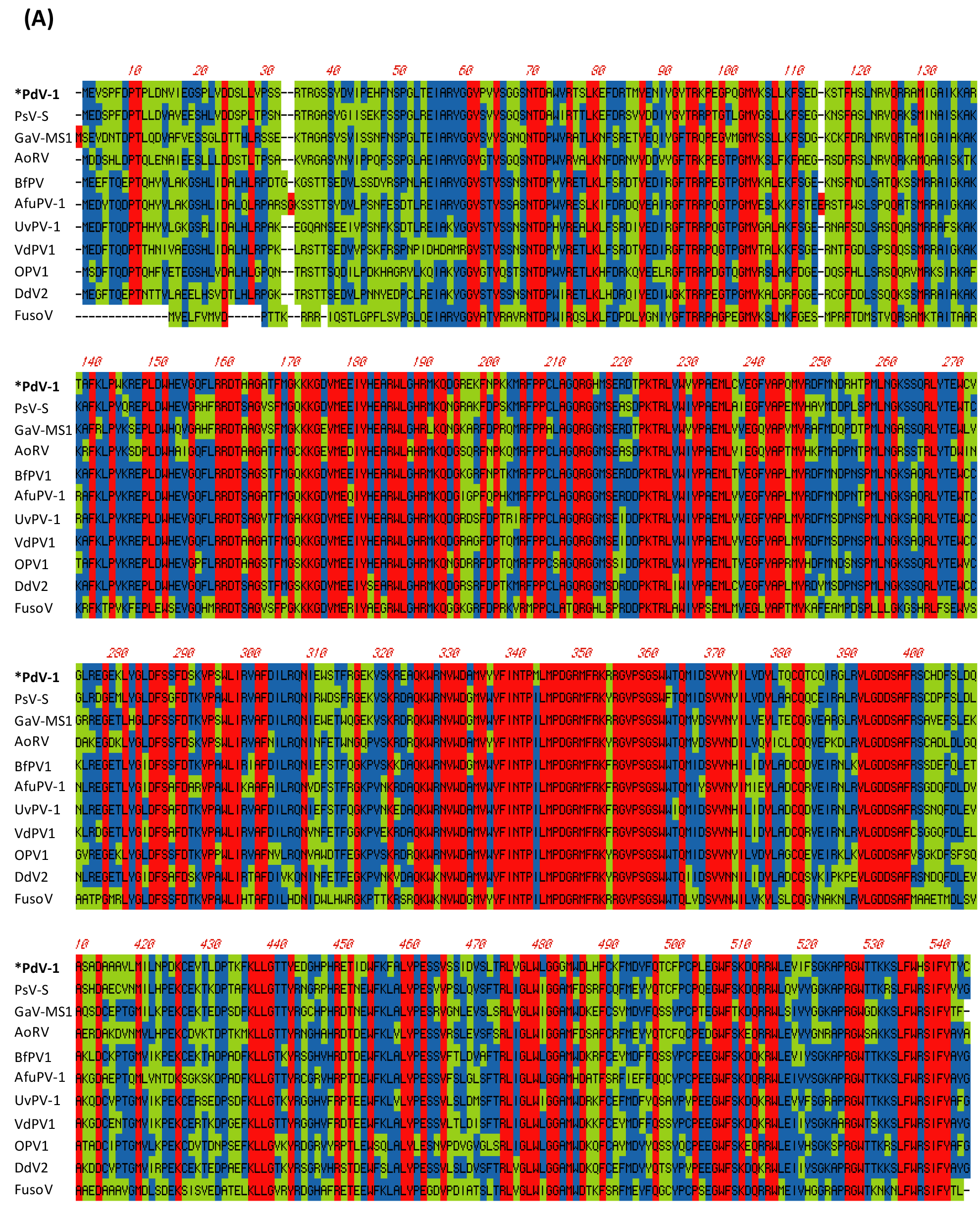
Comparison of the amino acid sequences of putative RdRp of the *Pseudogymnoascus destructans* virus (PdPV-1), *Penicillium stoloniferum* virus S (PsV-S), *Gremmeniella abietina* virus MS1 (GaV-MS1), *Aspergillus ochraceus* virus (AoV), *Botryotinia fuckeliana* partitivirus-1 (BfPV1), *Aspergillus fumigatus* partitivirus-1 (AfuPV-1), *Ustilaginoidea virens* partitivirus 1 (UvPV-1), *Verticillium dahliae* partitivirus 1 (VdPV1), *Ophiostoma partitivirus* (OPV1), *Discula destructiva* virus 2 (DdV2), and *Fusarium solani* virus 1 (FusoV). Red: 100% identity; Blue: consensus match; Green: mismatch.

**Fig. S2.**
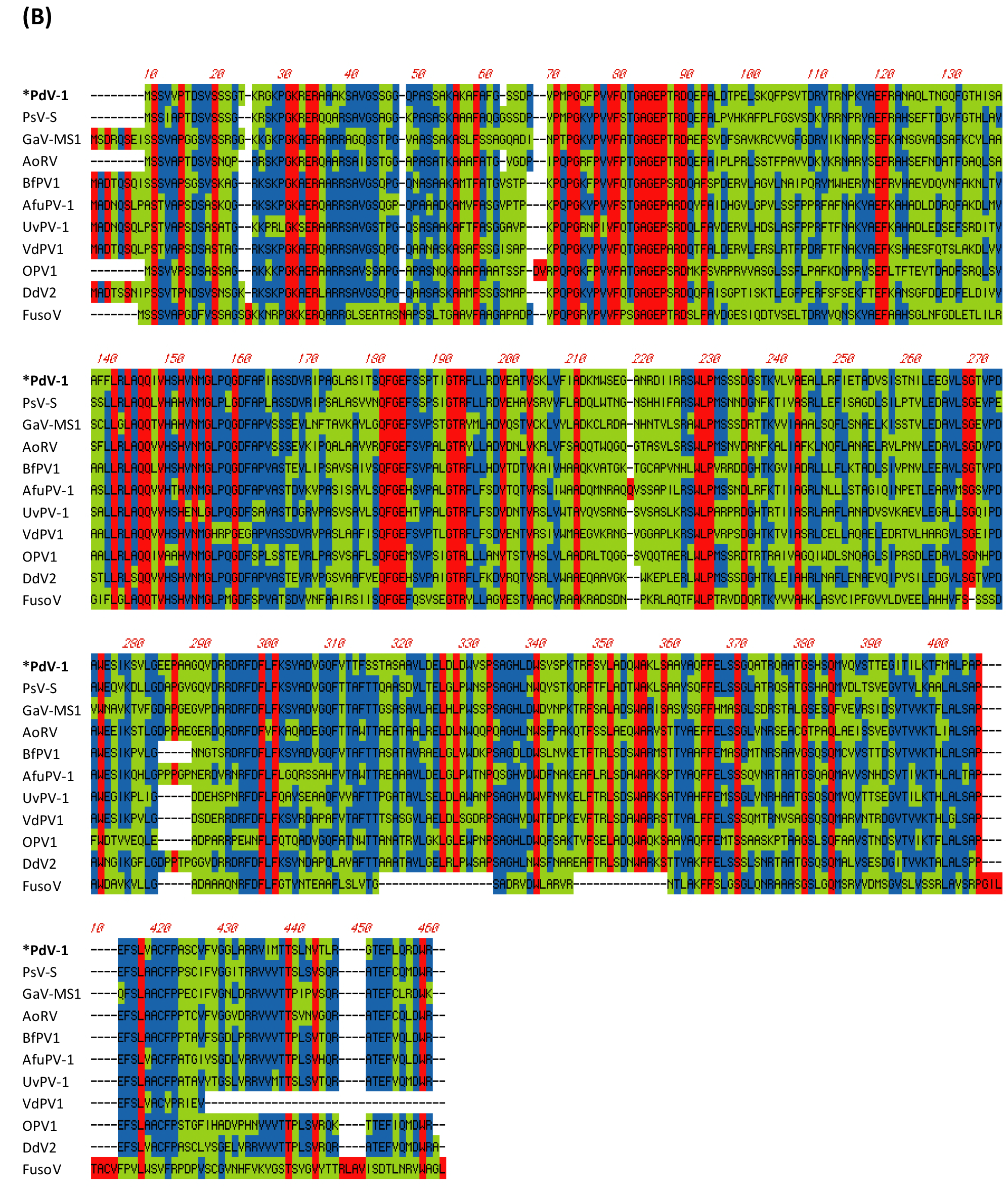
Comparison of the amino acid sequences of the capsid protein (CP) of the *Pseudogymnoascus destructans* virus (PdPV-1), *Penicillium stoloniferum* virus S (PsV-S), *Gremmeniella abietina* virus MS1 (GaV-MS1), *Aspergillus ochraceus* virus (AoV), *Botryotinia fuckeliana* partitivirus-1 (BfPV1), *Aspergillus fumigatus* partitivirus-1 (AfuPV-1), *Ustilaginoidea virens* partitivirus 1 (UvPV-1), *Verticillium dahliae* partitivirus 1 (VdPV1), *Ophiostoma partitivirus* (OPV1), *Discula destructiva* virus 2 (DdV2), and *Fusarium solani* virus 1 (FusoV). Red: 100% identity; Blue: consensus match; Green: mismatch.

**Fig. S3.**
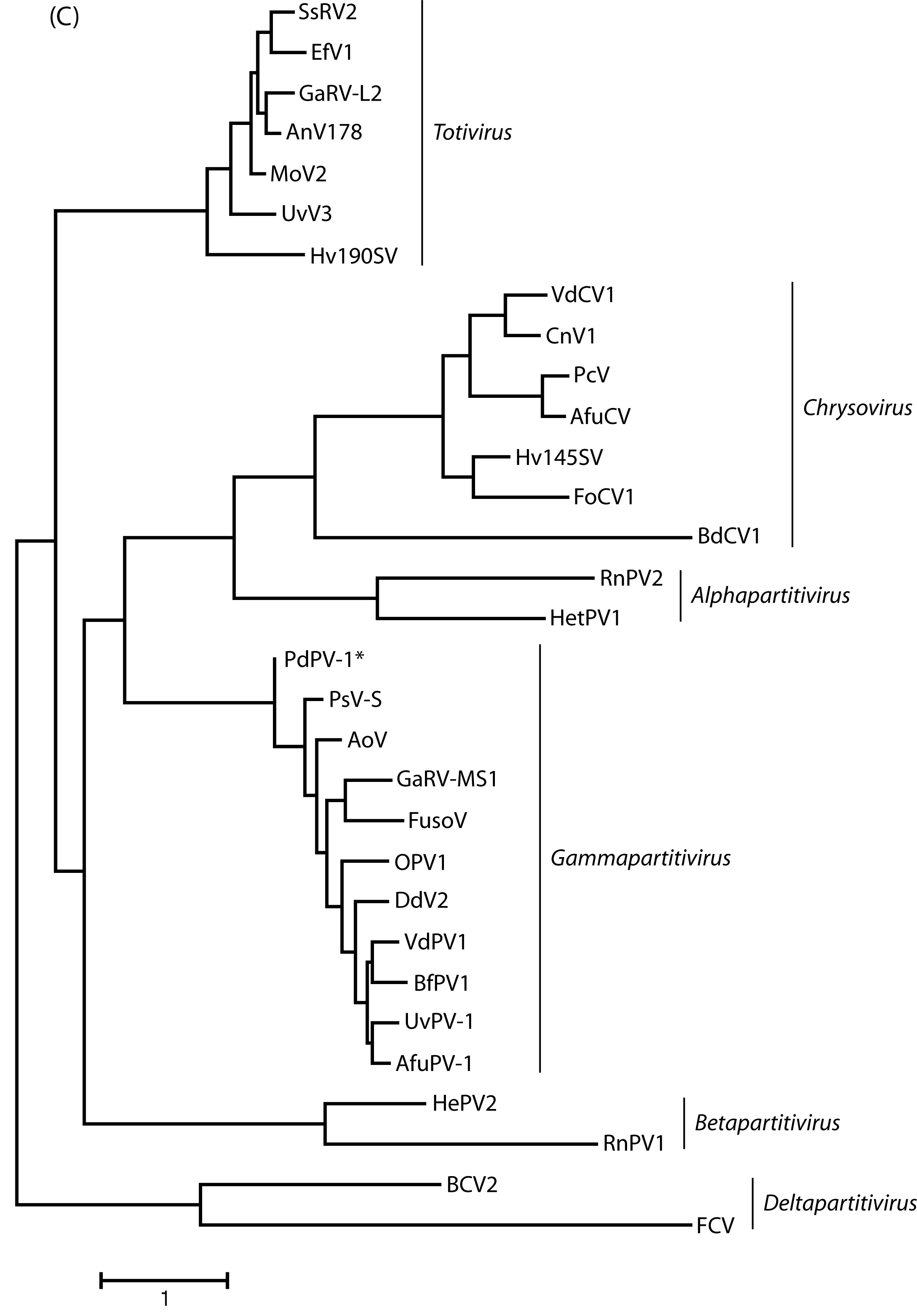
Phylogenetic analyses of PdPV-1. CP amino acid sequences of representative members of the family Partitiviridae, Totiviridae and Chrysoviridae were used to construct Maximum likelihood phylogenetic tree by MEGA 6 (GenBank accession numbers are provided in Table S2).

